# Cancer cell deformability impacts the rate of confined migration but not decision making

**DOI:** 10.1101/2025.05.12.653524

**Authors:** A. van der Net, R. C. Boot, I. van Dijk, J.P. Conboy, P.E. Boukany, G.H. Koenderink

## Abstract

Cancer cells can utilize different invasion strategies to overcome physical arrest during confined migration through tissues with small pores. Cancer cell plasticity allows switches between different migration modes and transitions between single-cell and collective migration. The biophysical parameters that guide these decisions are poorly understood. In this work we investigated the link between cell deformability and migration efficacy in constrictions of two mesenchymal cancer cell types with similar invasion strategies: HT1080 fibrosarcoma cells and MV3 melanoma cells. To this end, we designed microfluidic platforms for (1) high-throughput cell deformability measurements and (2) migration through a variety of confining geometries. We measured different deformabilities for HT1080 and MV3 cells and correlated this to their migration efficacy through confinements. However, higher deformability and improved squeezing ability did not impact decision-making at junctions of channels of different widths. Our findings show that cell deformability correlates with better squeezing abilities through confinements, but does not impact directionality decisions.

## INTRODUCTION

Invasion of tumor cells from a primary tumor site into the surrounding tissue is the first step of metastasis. Next, invasive tumor cells can intravasate into blood or lymph vessels, where they can travel to other organs and proliferate into new tumors. As metastasis accounts for over 90% of the mortality rates from cancer, understanding how cancer cells successfully invade local tissue with complex compositions is crucial to develop effective therapies^1–3^. Interestingly, despite being so lethal, tissue invasion is a highly inefficient process where most cells die from mechanical or oxidative stress or from successful attacks by our immune system^4,5^. Mechanical stress arises from cells invading into dense tissues, where small pore sizes force cells to deform and ‘squeeze’. During squeezing, contractile actin bundles confine the nucleus, which can lead to ruptures of highly deformed regions of the nuclear envelope and eventually loss of nuclear compartmentalization and DNA damage^6–8^. Therefore, single cell migration is limited by extracellular matrix (ECM) density and nuclear deformability, which can lead to physical arrest of cancer invasion when the pore size is smaller than 10% of the nuclear diameter^9^. However, cancer cells are highly adaptive and have different strategies to avoid physical arrest. One strategy is to deploy proteolytic enzymes such as matrix metalloproteases in order to degrade the matrix and create a migration path^9,10^. Another strategy is to switch to alternative motility mechanisms to better propel the nucleus through confinements, for instance switching to amoeboid or nuclear piston mechanisms^9,11–13^. Alternatively, mesenchymal cells can switch to collective migration in response to confinement^14,15^ to circumvent physical arrest in narrow pores^16^. Although single cells migrate faster, migrating cell clusters have more directional efficiency and result in more metastatic site formations^17,18^.

Although there is abundant evidence for cancer cell plasticity in response to confinement, it remains unclear how cells make the decisions between different invasion strategies to avoid physical arrest. An interesting side-by-side comparison was made in a study of Haeger et al.^16^, which compared the ability of human melanoma cells (MV3) and fibrosarcoma cells (HT1080), two mesenchymal cancer cell lines, to invade 3D collagen networks with pore sizes ranging from 35 µm^2^ to 3.5 µm^2^. Both cell types underwent a similar single-to-collective transition when the pore size was smaller than a threshold value around 16 µm^2^, a pore size in which cells have to start to deform to ‘squeeze’ through. Although HT1080 and MV3 cells deployed a similar strategy in response to physical confinement, HT1080 cell invasion was less dependent on matrix degradation and higher numbers of single HT1080 cells could still invade for every pore size that was tested. HT1080 invasion into small pores was hence less dependent on the individual-to-collective transition and on proteolytic strategies^16^.

We hypothesize that the greater ability of HT1080 cells to squeeze through narrow confinements compared to MV3 cells stems from a difference in cell deformability. To test this hypothesis, we studied how HT1080 and MV3 cell deformability is related to migration efficacy under confinement. In addition, we tested the relation of cell deformability to decision making when cells face a choice between channels of different widths. This was motivated by a recent study showing that cells migrate in the direction of least confinement to minimize energetic costs directed by force generation necessary to migrate through confining channels^19^. Although collagen-based hydrogels are a convenient reductionist model system to mimic the confinement imposed by the tumor microenvironment, their heterogeneous structure and wide distribution of pore spaces and connectivities make it challenging to interpret the exact effects of confinement on invasion strategies. Therefore, in this work we used microfluidic devices with standardized well-controlled pore spaces to analyze the effects of constriction. Another important benefit of this model is that the pore spaces are fixed, whereas cancer cells actively remodel collagen networks by mechanical forces and proteolytic degradation.

Here, we correlated measurements of cell deformability and confined migration for HT1080 and MV3 cells. We first designed a cell deformability microfluidic device to perform high-throughput cell deformability measurements, combining the capillary constriction design by Au et al.^20^, combined with the T-junction design from^21^. We could thus investigate how the deformability differs between the HT1080 and MV3 cell types when forced through vessel-sized constrictions by a flow. We subsequently studied the migration efficacy and decision making of the same cells in different confining geometries by introducing custom-made microfluidic migration devices, inspired by Davidson et al.^22^. Using the cell deformability device, we discovered that the HT1080 cells were more deformable than MV3 cells, especially for larger cell sizes (> 20 µm), where deformability was consistently larger than for smaller cells. Using the microfluidic migration devices, we found that HT1080 cells were better at crossing constrictions compared to MV3 cells, with a larger fraction of cells crossing (78% vs. 52%, respectively) and also crossing faster. Despite these differences in deformability and squeezing ability, decision making in narrow bifurcations was similar for HT1080 and MV3 cells. Both cell types showed a preference for wider constrictions, however, this preference was considerably reduced in narrower bifurcations. These results show that cell deformability correlates with better squeezing abilities through confinements, but does not impact directionality decisions.

## RESULTS

### HT1080 cells are more deformable than MV3 cells

Microfluidic devices mimicking capillaries from the blood vascular network were developed to assess cell deformability, consisting of 18 parallel microchannels narrowing into 10 × 10 µm^2^ constrictions (see schematic in Fig. 1A). Cells first flow through a large 100 × 100 µm^2^ main channel due to a pressure difference of 1 kPa between the cell entry port *P*_1_ and the perfusion port *P*_2_. They are then directed towards the constriction microchannels by a pressure gradient between the main channel and the outlet port *P*_3_, which is at atmospheric pressure. The two pressure inlets *P*_1_ and *P*_2_ allow precise control of the flow velocity of the cells through the main channel and the pressure forcing the cells into the microchannels. 3D COMSOL simulations of the device showed that the horizontal pressure gradient across the parallel microchannel array is negligible (∼30 Pa, Supplemental Fig. S2). They furthermore showed that the pressure that cells experience when entering the constrictions depends on whether the other channels are clogged by traversing cells. The pressure changed by ∼20% between the case of all channels being open (2.9 kPa) versus closed (3.5 kPa, Supplemental Fig. 9B). While this pressure change is expected to influence the entry and traversal time of cells crossing the constrictions^20^, it does not influence the cell deformability we focus on in this work.

**FIG. 1.**
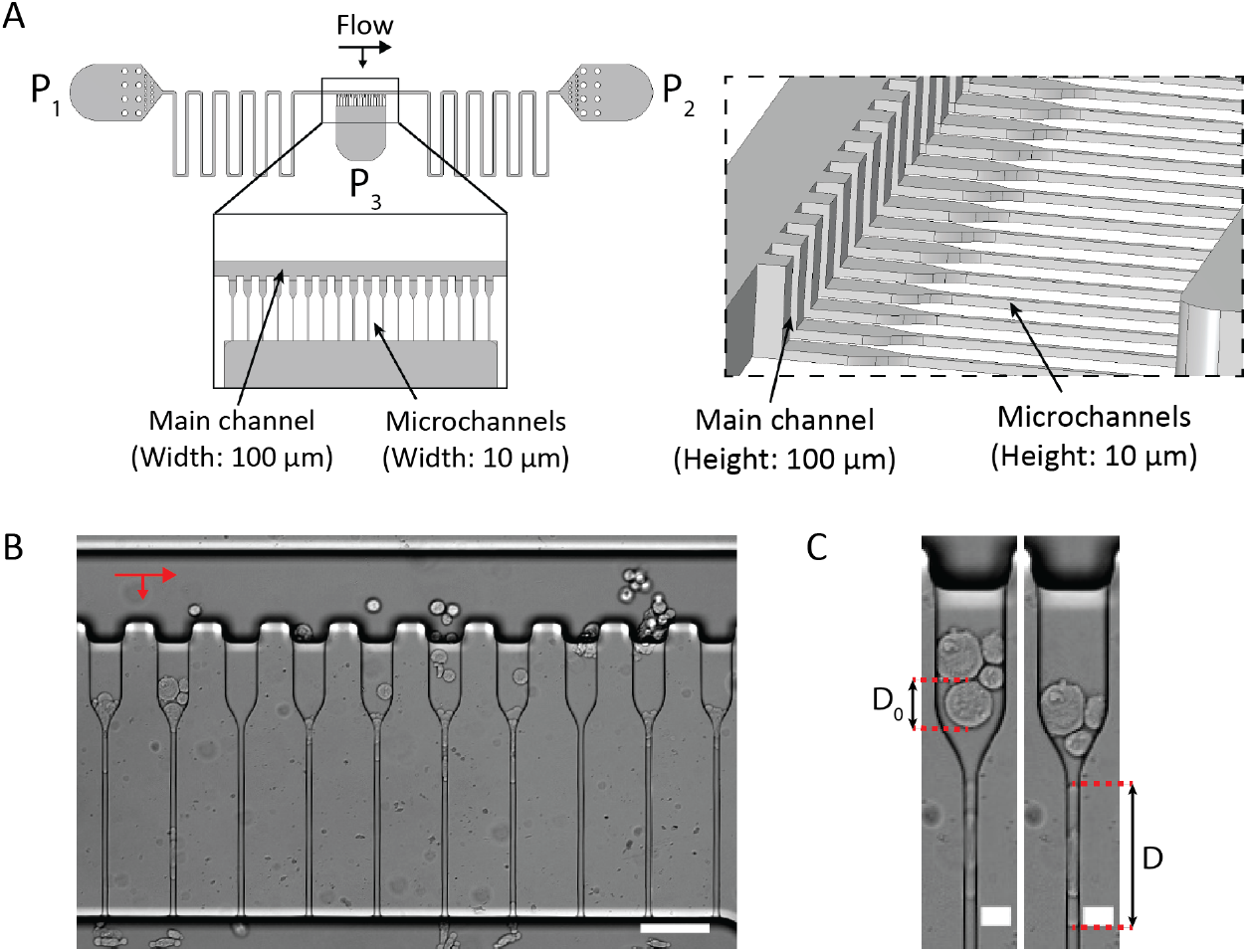
Overview of the microfluidic cell deformability device. (A) Schematic showing (left) the top view of the design together with a close-up top view of the main channel and the parallel constriction microchannels, and (right) a tilted side view showing the differing heights for the main channel and the microchannels. (B) Bright-field top view image of the parallel microchannels with HT1080 cells flowing freely through the main channel and being squeezed through the microchannels due to a pressure gradient from top to bottom. Red arrows indicate the flow directions. Scale bar is 100 µm. (C) Bright-field images of a HT1080 cell entering a constriction inlet with a diameter *D*_0_ (left panel) and subsequently stretching in the microchannel to a length *D*. Scale bars are 20 µm.

At the inlet of the 50 µm-wide microchannels, cells are only confined in height, to a diameter *D*_0_. When the cells enter the 10 µm-wide constrictions, they are also confined in width, leading to a deformation length *D* (Fig. 1B-C). We determined the dimensionless cell strain from *D*_0_ and *D* using Eq. (1). As shown in Fig. 2, the strain increased with cell size (*D*_0_) for both HT1080 and MV3 cells. The larger the cell, the more it has to stretch in the channel due to the confinement. We therefore binned the cells by sequential pairs of *D*_0_ (e.g. cells with *D*_0_ = 15 µm and 16 µm were combined in one bin), and compared *S* between the respective bins for each cell type. This analysis revealed that the HT1080 cells consistently deformed more than the MV3 cells for the same cell size (note that the two cell types had similar size distributions S8). These findings suggest that cell volume is not conserved, as cells with the same original volume in the constriction inlet display different deformations. Using Eq. (5), we determine a theoretical strain *S*_∗_ in case of volume conservation, which is indicated in Fig. 2 by the red dotted lines. Comparing this theoretical limit to the data, we find that both the MV3 and HT1080 cells increased in volume in the constrictions.

**FIG. 2.**
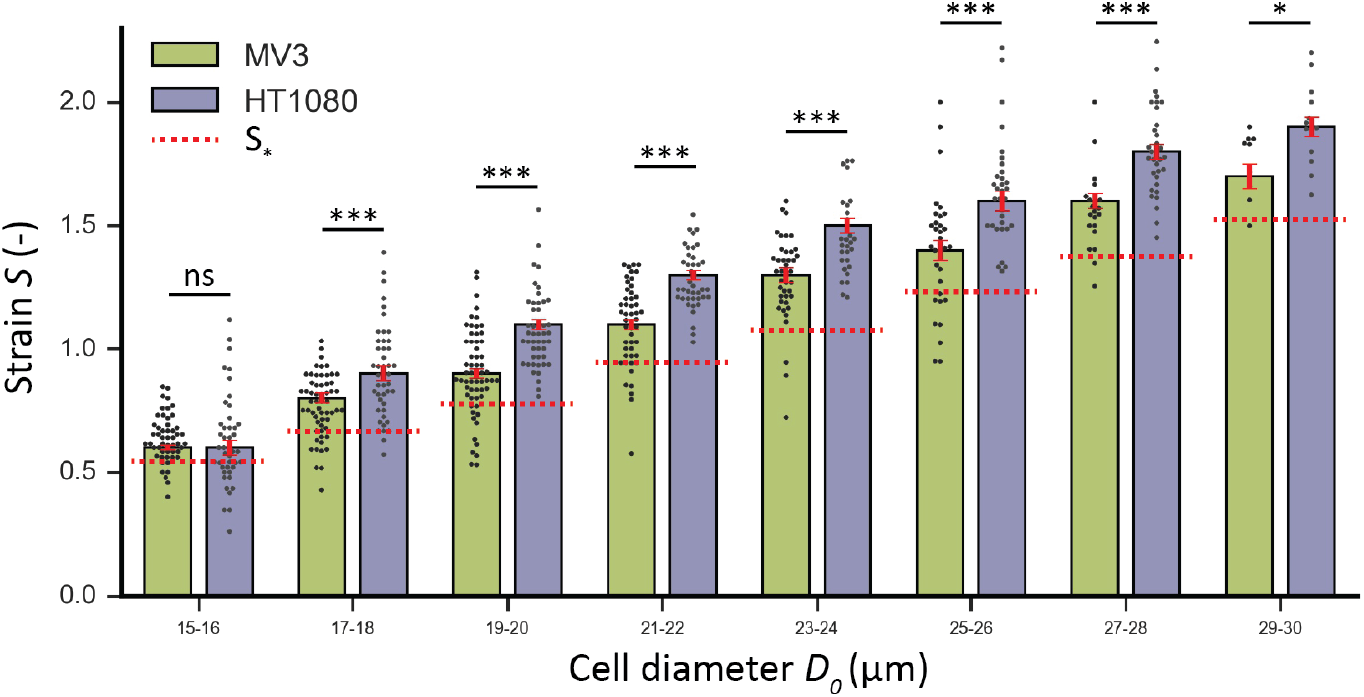
Histograms comparing the strain *S* for MV3 and HT1080 cells measured using the microfluidic cell deformability device. Cell diameters *D*_0_ were binned for MV3 (green) and HT1080 (purple) cells. Red dotted lines depict the theoretical strain *S*_∗_ in the limit of volume conservation, calculated using Eq. (5) and inserting the largest *D*_0_ for each bin (e.g. *D*_0_ = 16 µm for the 15-16 bin). (*) = p < 0.05, (***) = p <0.001 (n.s.) = nonsignificant. Error bars are SEM. N = 300 cells per condition.

### Microfluidic cell migration device design

To test whether the difference in HT1080 versus MV3 cell deformability affects their ability to migrate through narrow constrictions, we made three different microfluidic designs based on previous work by Davidson et al.^22^. All three designs have two media reservoirs on both sides of the device, connected with a wide bypass channel to enable fast equilibration of fluid height between both reservoirs (Fig. 3A). Furthermore, they all have a constriction area with parallel migration channels that were functionalized with type I collagen to promote cell adhesion and crawling. Instead of long, straight channels, we designed pillars or walls made of interconnected arcs, to mimic the discontinuous spatial environment found in the interstitial matrix of connective tissues (Fig. 3B). Moreover, we chose constriction widths in the range of 5 µm to 30 µm, to mimic the range of pore sizes found in tumor microenvironments^23^. The first design had an array of parallel channels with sequential constrictions of 5 µm. The second design had Y-junctions that start as 30 µm wide channels and then split into either a 20 µm and 10 µm wide channel, or into a 10 µm and 5 µm channel. This design allowed us to investigate how cell deformability influences the migration path when cells face a choice between different constrictions.

**FIG. 3.**
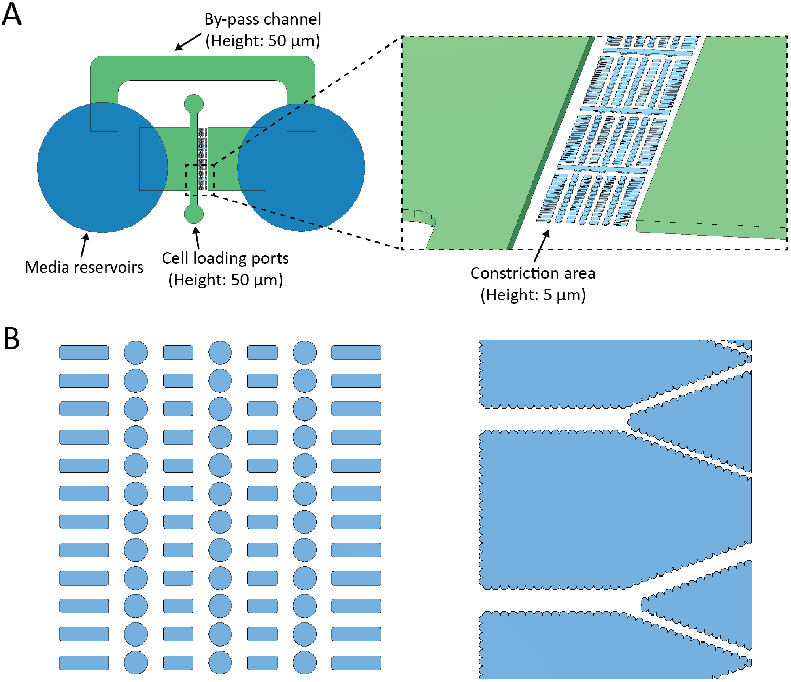
Overview of the microfluidic cell migration device. (A) Left: Schematic showing the top view of the design. The media reservoirs are punched (dark blue) to connect the by-pass channel and cell loading ports (green). Right: a tilted side view showing the constriction area between two non-constricted chambers. (B) The two different designs for the constriction areas: (left) rows with identical constrictions of 5 µm in width between the circular pillars, and (right) Y-junctions with a 30 µm wide channel, lined with inverted catenaries, splitting with a 40° angle into either a 10 µm and 5 µm wide channel (above), or into a 20 µm and 10 µm wide channel (below). Note that the pillars and the walls made of interconnected arcs mimic the discontinuous spatial environment that cells encounter when invading interstitial matrices.

### HT1080 squeeze faster through constrictions than MV3 cells

To determine if higher deformability is associated with faster migration of cells through narrow constrictions, we performed migration experiments for HT1080 and MV3 cells in the microfluidic migration device with identical constrictions of 5 µm. From the time-lapse image series, we identified all squeezing cells and measured the time duration spent by cells and their nuclei in the constriction areas, defined as the rectangle between the diameter of two adjacent pillars (Fig. 4A-C, see S3 for links to full videos; multimedia available online). Migrating cells were defined as cells that entered a constriction area during imaging and exited the constriction area on the opposite side during imaging. Cells were classified as stuck when they spent > 10 frames (equal to 2 hours) in the same constriction area without exiting during imaging. We found that with the same seeding densities, HT1080 cells had a larger number of cells squeezing through the constrictions compared to MV3 cells, while they had a similar amount of cells stuck in a constriction (Fig. 4D). In addition, HT1080 cells more often squeezed multiple times (2 or 3 times) during the time frame of imaging (around 16 hours) compared to MV3 cells, which on average only migrated through one constriction during imaging. To determine the amount of time it takes for cells (Fig. 4E) and their nuclei (Fig. 4F) to squeeze through a constriction, we measured the number of frames with a Cytotracker (cytoplasm) or Hoechst (nucleus) signal. For both cells and nuclei, HT1080 cells were significantly faster at squeezing compared to MV3 cells. After squeezing through a constriction, cells often showed nuclear deformations and/or ruptures, often also leading to migration inhibition and/or cell death. These deformations were more prominent in MV3 cells than in HT1080 cells (Fig. S4). Altogether, these results show that HT1080 cells, with a higher deformability, are indeed more efficient at squeezing through narrow pores as compared to MV3 cells.

**FIG. 4.**
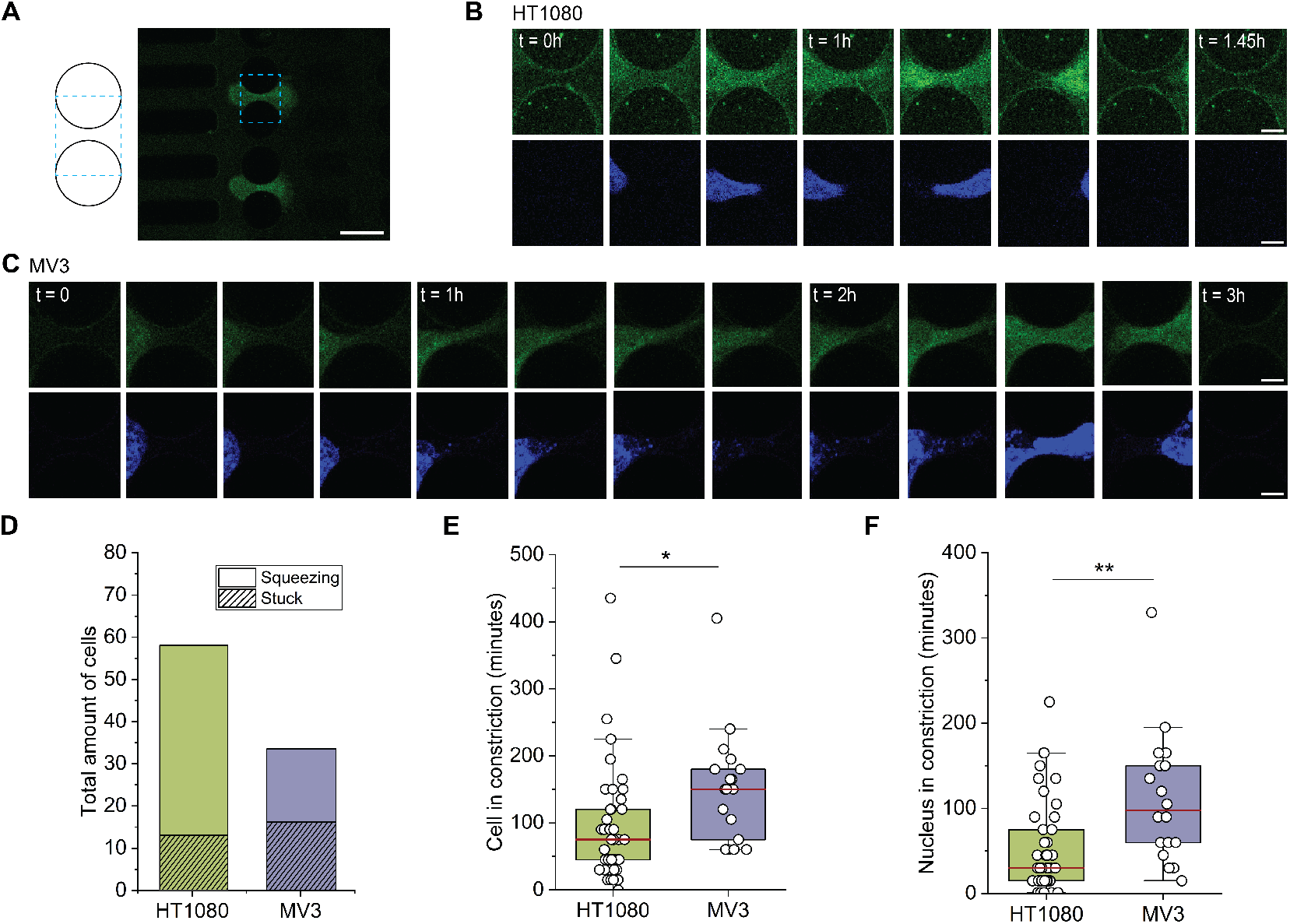
Analysis of confined cell migration efficacy in arrays of pillars. (A) Schematic and example fluorescence image showing how the constriction region was defined as the rectangular area (dashed box) formed by connecting the diameters of two adjacent pillars (circles). Scale bar is 15 µm. (B,C) Time lapse fluorescence image series showing HT1080 cells (B) and MV3 cells (C) squeezing through constrictions. The images are cropped to show the constriction region and show the cell cytoplasm labeled with Cytotracker Orange in green and the nucleus labeled with Hoechst 33342 in blue. Scale bars are 5 µm. (D) Stacked bar graph showing the total number of cells in constriction regions for HT1080 (green) and MV3 (purple) cells, separated between cells that actively squeezed through the constrictions (smooth) and cells that never squeezed through the area over an observation time of 13-14 hours (striped). (E,F) Boxplots for the amount of time (in minutes) that cells and nuclei were detected in the constriction areas, derived from the number of frames with 15 minutes time intervals. (*) = p < 0.05, (**) = p < 0.005.

### HT1080 and MV3 cells make similar migration decisions in narrow bifurcations

To understand if the higher deformability and squeezing efficiency of HT1080 cells compared to MV3 cells also impact directional decisions at junctions between channels of different widths, we next performed migration experiments in the microfluidic migration chips with bifurcation designs. In these chips, both cell types were subjected to constriction designs of 30 µm wide channels splitting into either 20 µm- and 10 µm-wide channels, or into 10 µm- and 5 µm-wide channels. To analyze cellular decision making, we performed live-cell imaging of the constrictions to visualize the migrating cells (Fig. 5A). We then analyzed which size channel the cells migrated into from the wider channel (Fig. 5B). For the 20 µm vs. 10 µm bifurcation, both cells had a comparable preference for the wider (20 µm) channel, with 69.2% (HT1080) and 72.7% (MV3). Interestingly, when presented with smaller constrictions of 10 µm vs. 5 µm, both cell types lose their clear preference for the larger constriction, with 52.6% (HT1080) and 58.3% (MV3) of cells migrating into 10 µm channels. These results show that HT1080 and MV3 make very similar decisions when presented with confinements of different sizes.

**FIG. 5.**
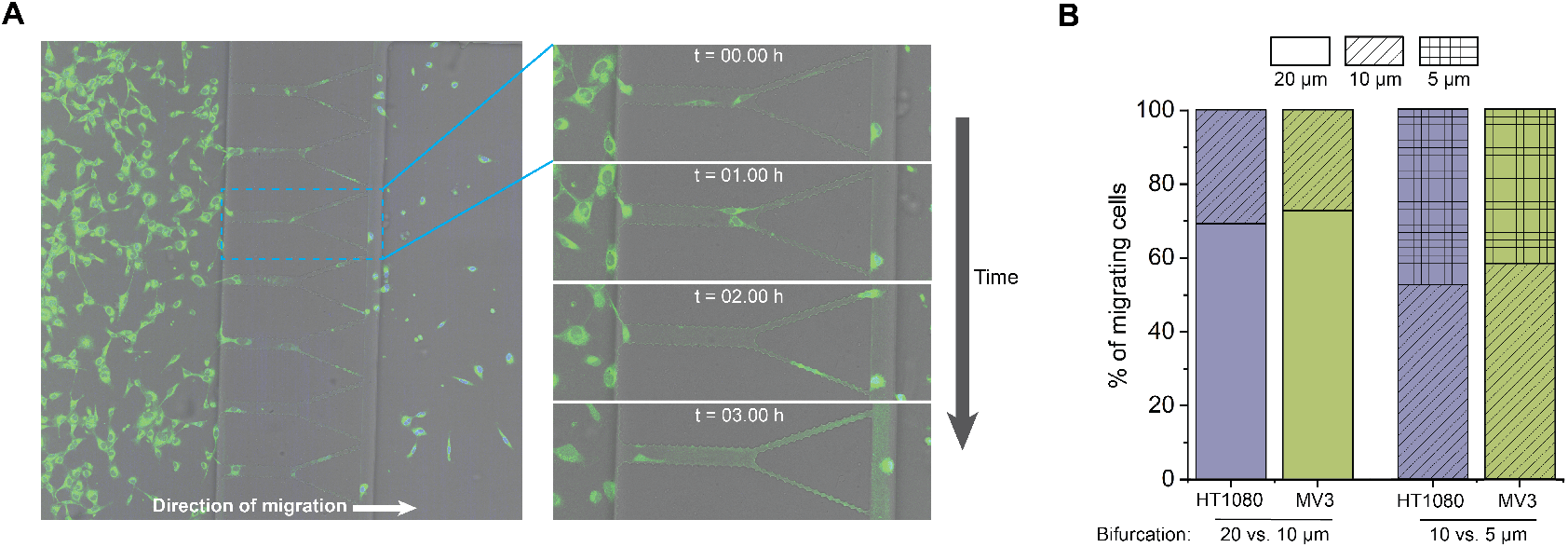
Decision making of migrating cells at narrow bifurcations. (A) Overview of microfluidic chip in which cells (green) migrate through 30 µm channels with a bifurcation of 5 & 10 µm. Blue square indicates zoomed-in area at 4 different time points, showing two cells migrating. (B) Stacked bar graph showing the decision making of cells at the bifurcations, analyzed by visual inspection and converted to percentages, for bifurcations of 20 µm (smooth) versus 10 µm channels (striped) and of 10 µm versus 5 µm (cross-hatched).

## DISCUSSION AND CONCLUSION

The goal of this work was to test whether the different strategies of HT1080 and MV3 cells to perform confined migration correlate with their deformabilities. Our work was inspired by a study by Haeger et al. (2014), which showed that, even though both cells have similar mesenchymal features, HT1080 cells are able to squeeze through smaller pores than MV3 cells without resorting to MMP-mediated matrix degradation or switching to collective invasion^16^. To compare the mechanical properties of the two cell types, we first designed a microfluidic device with parallel constriction channels for high-throughput cell deformability measurements. Using this device, we found that HT1080 cells deformed more in the channels than MV3 cells when comparing cells of similar sizes. We also found that both cell types exhibit a change in volume when forced into the narrow constrictions. A recent study reporting high-speed morphological measurements on cells migrating into narrow constrictions also showed changes in cell volume^24^. However, the exact process behind this dynamic volume change is not clear. Likely, plasma membrane channels and transporters, which transport intracellular osmolytes (inorganic ions and small organic compounds) in and out of the cell^25,26^, are involved. These transport processes generate osmotic gradients, which drive water across the cell membrane through aquaporin water channels^27^. Directed water permeation has indeed been implicated in confined cell migration, where polarized distributions of Na^+^/H^+^ transporters and aquaporins facilitate volume changes^28^. To test this mechanism, it will be interesting to measure the change in cell deformability in presence of inhibitors of membrane transporters and aquaporins to manipulate or block water exchange^29^.

To test whether the cytoskeleton plays any role in confinement-induced cell deformability, we treated MV3 cells with cytochalasin D to depolymerize actin and measured their deformability in the constrictions. Microindentation experiments showed an average ∼5-fold decrease of the effective Young’s modulus of the cells (Fig. S5), confirming successful inhibition of actin polymerization^30^. By contrast, the deformability of the MV3 cells in the microfluidic constrictions was completely unaffected by the CD treatment (Fig. S6). This suggests that, at least for these cells, the actin cytoskeleton does not affect cell deformability. We note that previous work comparing MCF7 and MCF10 breast cancer cells did suggest a correlation between cell deformability in constrictions and cellular elasticity^31^. Therefore it will be interesting to test the role of actin in cell deformability further, together with the role of other cytoskeletal components, especially the vimentin cytoskeleton. Actin and vimentin form interpenetrating networks and jointly determine cell deformation under compressive and tensile loading^32,33^. Western Blot analysis shows that the HT1080 cells express higher levels of vimentin than the MV3 cells (Fig. S7), which could potentially enable them to deform more without damage^12,34,35^. Future experiments with vimentin-knockout cells could provide insight in the contribution of vimentin to cell deformability.

Other intermediate filaments of interest are nuclear lamins, which govern nuclear deformability and thereby impact confined migration^9,36^. Nuclear stiffness is mainly dependent on the levels of lamin A/C^37^, and recent work by Amiri et al.^38^ demonstrates that nuclear lamins respond similarly to deformations during both aspiration and active cell migration. Since the nucleus is the limiting factor in cell passage through constrictions, it will be interesting to quantify the lamin A/C expression levels of the HT1080 and MV3 cells as well as the size of their nuclei.

To couple the deformability of the cells to their ability to migrate through constrictions that are narrow enough to cause physical arrest, we next designed a second microfluidic device that allows for live-cell imaging of migration through a 3D environment of narrow constrictions with varying sizes. Here, we found that HT1080 could migrate faster through small constrictions compared to MV3 cells, with fewer cells getting stuck. Physical arrest of cell invasion is known to be controlled by cellular deformability and extracellular confinement^9^. However, studies on cancer cell invasion in different confining microenvironments, including extracellular matrix gels and microfluidic constrictions, have shown that cancer cell plasticity can prevent physical arrest and enhance the invasion potential. Cancer cells can adopt different strategies including migration mode switching^9,11–13^, proteolysis of the extracellular matrix^9,10^, and individual-to-collective migration transitions^14–16^. However, the decision-making process behind these choices is not well understood. There is evidence that decisions between different strategies are interdependent; both mesenchymal-amoeboid migration mode switching and individual-to-collective transitions in response to confinement depend on proteolytic activity^16,39,40^. Additionally, collective invasion can also directly transition to single-cell amoeboid migration in response to hypoxia^41^. A side-by-side comparison between HT1080 and MV3 cells by Haeger et al.^16^ showed that these cell types have similar invasion strategies but with subtle differences: HT1080 cells were less dependent on matrix proteolysis to squeeze through collagen gels with narrow pores. Our comparison of the single-cell migration speeds of these cells through microfluidic constrictions confirmed that HT1080 cells can squeeze faster through narrow pores and get less easily stuck. Moreover, we showed that this difference correlates with higher cell deformability. This correlation is consistent with earlier findings, where cell deformability was coupled to the metastatic potential of cancer cells^42–44^.

Finally, to understand how cell deformability and migration efficacy impact cellular decision making in dense environments, we performed migration assays using bifurcation designs in the same migration microfluidic set-up. We found that decision making was similar for HT1080 and MV3 cells and thus not influenced by their deformabilities nor migration efficacies. Cells showed a preference for larger constrictions when presented with a decision between 20 µm and 10 µm constrictions, but this preference was lost when presented with narrower constrictions of 10 µm and 5 µm. Similar experiments with constriction bifurcations by Zanotelli et al.^19^ described decision-making to be mainly determined by cell stiffness (Young’s modulus). This finding suggests that HT1080 and MV3 cells have a comparable Young’s modulus. Indeed microindentation experiments confirmed that the cells had a comparable effective Young’s modulus of ∼500 Pa (Fig. S5). Although migrating cells generally prefer wider constrictions to reduce energetic costs^19,45^, both HT1080 and MV3 cells lost their clear preference for wider constrictions in the bifurcations of 10 µm versus 5 µm. Because the bifurcations designs in Zanotelli et al.^19^ had similar constriction sizes (12 µm versus 7 µm) and they did not report any loss of preference, we hypothesize that energy costs might plateau under a certain constriction size. This size might potentially depend on cell and nucleus size; for our cells, it is around ∼10 µm.

Gaining a deeper understanding of how the physical parameters of cancer cells relate to migration efficacy, invasion strategies and therapy resistance can be of great value for clinical applications. Computational modeling based on spheroid-collagen invasion assays showed that tumor invasiveness is maximized by heterogeneity in cell deformability and cell size^46^. Furthermore, deformability heterogeneity could potentially also promote collective invasion strategies by enabling cells to dynamically adjust their position within the cluster^46^. As recent work also revealed that collective invasion *in vivo* can create radiation-resistant niches^47^, it is important to further study how cell mechanics contribute to this invasive strategy. Moreover, recent studies prove the predictive power of deformability measurements of patient-specific cancer cells for the efficiency of different treatments, indicating the potential for the development of new prognostic tools^30,48^.

## METHODS

### Cell culture

Human fibrosarcoma HT1080 cells (ATCC, CCL-121) and human melanoma MV3 cells were gifted by Peter Friedl (RadboudUMC, the Netherlands). The HT1080 cells are human male fibrosarcoma cells derived from biopsy, as described in ref.^49^. The MV3 cells are human male melanoma cells that were three times xenografted in nude mice and selected for highly metastatic behavior as described in ref.^50^. HT1080 cells were cultured in High Glucose Dulbecco’s Modified Eagle’s Medium (DMEM, #11574486, Thermo Fisher) and MV3 cells were cultured in DMEM/F12 1:1 medium (#11520396, Thermo Fisher), both supplemented with 10% Fetal Bovine Serum (FBS, Gibco) and 1% penicillin-streptomycin (Sigma-Aldrich). Cells were incubated at 37 ° C and 5% CO_2_ and subcultured at 80-90% confluency, with regular tests for mycoplasma infections. Cell counting and cell size measurements were performed using a Countess 3 FL Automated Cell Counter (Thermo Scientific).

### Actin polymerization inhibition

Cells were incubated for a minimum of 30 minutes with culture media supplemented with 50 µM Cytochalasin D (Merck Sigma, #C2618) or 0.025 µL/mL DMSO (Bioke, #12611P) as a control. Previous research has shown that incubation of fibroblasts with this concentration of CD reduced the cell stiffness as measured by uniaxial stress-strain testing by 50 %^51^.

### Cell deformability device fabrication

Cell deformations were tested in a custom-designed microfluidic device that combines aspects of devices reported by Au et al.^20^ and by Davidson et al.^21^. The design is available in DWG format at https://github.com/RubenBoot/CellandClusterDeformation). By fusing the parallel microchannel array from^20^ with the T-junction design from^21^, which incorporates a main channel that leads past the array, the overall volumetric flow rate through the device was greatly increased compared to the original design by Au et al.^20^ thus improving the throughput of cells reaching the microchannel array and traveling through the constrictions. The microchannels had dimensions of 10×10×280 µm^3^ (width by height by length), while the main serpentine channel was 100 µm wide and high. This multi-layered design was created using standard soft lithography at the Kavli Nanolab Delft, with a µMLA laserwriter (Heidelberg Instruments). For the first layer containing the microchannel array, SU-8 2010 photoresist (Kayaku Advanced Materials) was spun on a clean 4-inch silicon wafer to a height of 10 µm. The coated wafer was then baked at 95 °C for 3 minutes, after which the first layer was written and post-baked at 95 °C for 4 minutes. After development with SU-8 developer (Propylene glycol monomethyl ether acetate (PGMEA), Sigma-Aldrich), a second layer was spun to a height of 100 µm using SU-8 2050 (Kayaku Advanced Materials). It was now baked at 65 °C for 5 minutes followed by 95 °C for 15 minutes, written, post-baked at 65 °C for 3 minutes followed by 95 °C for 10 minutes and developed again. The heights of the SU-8 features were determined to be within ±10% of the desired heights using a Dektak Stylus Profiler (Bruker). Lastly, the master wafer was coated with trichloro(1H,1H,2H,2H-perfluorooctyl)silane (Sigma-Aldrich) to allow for easy demolding. The devices were made by pouring PDMS (Sylgard 184, Dow Corning) and curing agent at a mixing ratio of 10:1 (w/w) onto this master mold. The PDMS was degassed and cured at 65 °C for 3 hours. After curing, the PDMS was peeled off, 2 mm inlets and outlets were punched using a revolving punch plier (Knipex), and the devices alongside glass coverslips were plasma cleaned (Harrick Plasma) at 30 W for 2.5 minutes. The devices were kept in the oven at 65 °C to bond overnight.

### Cell deformability assays

The connective ports of the cell deformability device served for cell entry (*P*_1_), inlet of perfusion buffer (*P*_2_) and an outlet (*P*_3_) (Fig. 1A). The long serpentine main channel served to minimize the pressure drop over the parallel constriction channels at the center of the design. The devices were first filled and incubated for 45 minutes with 1% Pluronic F127 (Sigma) in phosphate buffered saline (PBS, Sigma), to decrease cell adhesion to the PDMS and glass surfaces. PTFE 008T16-030-200 tubing (Diba Industries, inner diameter 0.3 mm, outer diameter 1.6 mm) was cut into three pieces with an identical length of 50 cm (to prevent the influence of different pressure drops over tubing of differing lengths) and then flushed with the perfusion media using an MCFS-EZ pressure controller (Fluigent). After connecting each tubing piece to one of the connective ports, the Pluronic solution, PDMS debris particles and possible air bubbles were flushed out with cell culture medium by inducing a flow from *P*_2_ and *P*_3_ towards *P*_1_, with *P*1 connected to a waste tube. Once the main serpentine channel was free of obstacles, the tubing of *P*_1_ was connected to the cell suspension while a minor flow was still present from *P*_2_ to *P*_1_, to prevent air from entering the tubing when connecting the cell suspension. A cell suspension of 1 mL (with cell concentrations ranging between 0.5–3.5 × 10^6^ cells/mL depending on the experiment) was used. After reversing the flow direction, cells were subsequently flown through the main channel using a pressure gradient of 10 mbar (*P*_1_ at 40 mbar, *P*_2_ at 30 mbar). Outlet *P*_3_ was kept at atmospheric pressure, such that the cells felt the pressure difference near the center of the design and were forced into the constriction channels (Fig. 1B). To compare the deformability of MV3 cells and HT1080 cells, we performed experiments sequentially on the same deformability device. After collecting data for one cell type, the main channel was flushed with medium by connecting a tube of medium after reversing the flow from *P*_2_ to *P*_1_ to prevent air from entering the tubing at *P*_1_. Next, a suspension with the other cell type was connected to the device for further measurements.

Brightfield images of the deforming cells were captured using an inverted fluorescence microscope (Zeiss Axio Observer) with a 10x/NA 0.45 air objective and ORCA Flash 4.0 V2 (Hamamatsu) digital camera with a resolution of 2048×2048 pixels. We recorded time lapse image series of 30 seconds long, using an interval of 25 ms, with 13 constrictions fitting the field of view. Multiple sets were recorded per chip to capture data for a large number of cells, and the device was always discarded at the end of the experimental day. For all experiments, the cell suspension was kept at 37 °C using a Compact Dry Bath incubator (S 200-240V, Thermo Fisher Scientific) and time lapse image series were only recorded during the first hour after connecting the cell suspension to the chip, eliminating the need for CO_2_ injection.

### Passive transit image analysis

Analysis of the deformation of cells in the constrictions was conducted using Fiji^52^. For each time lapse image series, multiple cells squeezed through the parallel constriction channels. Only the cells for which both the original diameter *D*_0_ in front of the constriction (Fig. 1C, left panel) and the deformed length *D* once fully entered in the constriction (Fig. 1C, right panel) were clearly visible were included in the analysis. As the cells were not always perfectly round at the inlet, *D*_0_ was determined by taking the average of the long and short diameters of the cell. Using Fiji, *D*_0_ and the deformed length *D* for each cell were measured manually. Next we determined the strain *S* (dimensionless cell deformation) by taking the following ratio:

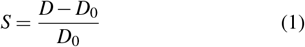

The theoretical strain *S*_∗_, being the strain expected in case of cell volume conservation, was calculated based on the cell volume estimated from the measurement of the cell size before it entered the constrictions. Assuming the cell in the inlet to be an approximate cylinder with a diameter *D*_0_ and a height *H* of 10 µm (Supplementary Fig. S1A), we know the volume *V*_1_ of a cell in the inlet to be:

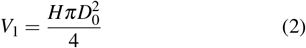

Once the cell enters the constriction channel, we approximate it as a beam of length *L* with a hemisphere on each end, having a radius *R* equal to half the width *W* and height *H* of the constriction (here, *R* = 5 µm) (Supplementary Fig. S1B). From this, the volume of the cell *V*_2_ in the constriction is found as:

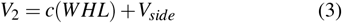

Here 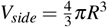 represents the combined volume of the two half-spheres. Assuming volume conservation (*V*_1_ = *V* 2), the length of the beam *L* is found by equating Eq. (2) and Eq. (3):

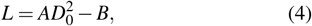

with *A* = *π/*4*W* and *B* = *V*_*side*_*/WH*. Inserting Eq. (4) into Eq. (1), we finally find the theoretical strain *S*_∗_ as:

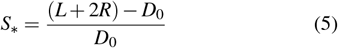

### Modeling of pressure distribution in the cell deformability device

Due to the T-junction design of the cell deformability device, a pressure gradient exists both horizontally and vertically along the constriction channel array. To determine the size of these gradients, we computationally modeled the pressure distribution inside the device using the finite element modeling software COMSOL Multiphysics 5.6. We considered the fluid flow in the 3D-model to be laminar and following the Navier-Stokes equation. The pressure gradient was examined for two extreme cases: (1) all constrictions are open, and (2) the flow through all constrictions is blocked (this occurs when all channels are clogged by traversing cells). The pressure drop over a tubular channel with laminar flow scales with the length of the channel and the inverse of the channel radius to the fourth power. As the tubing used in our experiments had a long length of 50 cm and a cross-sectional area with the same order of magnitude as the main channel, the pressure drop over the tubing was non-negligible. For this reason, the tubing was modeled by including 50 cm long rectangular channels with a 300 × 300 µm^2^ cross-section connected to all three ports. We used boundary conditions of 4 kPa at the far edge of the channel connecting to the cell entry port *P*_1_, 3 kPa at the edge of the channel connected to the perfusion port *P*_2_, and 0 kPa at the edge of the channel connected to the outlet port *P*_3_, similar to the pressures used in the experiments.

### Microindentation measurements

The effective Young’s modulus of the cells was measured using a Chiaro Nanoindenter (Optics11 Life), using an optical fiber probe with a stiffness of 0.027 N/m and a spherical tip with a radius of 3 µm. Measurements were performed on 14 cells per condition, attached to the glass bottom of a 35 mm dish (#81218-200, Ibidi) in CO_2_ independent medium (#11580536, Thermo Fisher). Cells were seeded at least 24 hours before experiments in order to adhere to the bottom. Indentations were made above the cell center with a loading rate of 2 µm/s. The modulus was calculated using the Hertzian contact model from the Optics 11 Life data viewer software (version 2)^53^. Measurements without a distinct contact point or with an otherwise unreliable model fit (*<* 0.9*R*^2^) were regarded as outliers and discarded from further analysis.

### Microfluidic migration device fabrication

We designed a microfluidic device tailored for observing cell deformation during migration through different custom-designed constriction areas inspired by Ref.^22^. The multi-layered master mold was created using standard soft lithography at the Kavli Nanolab Delft, with a µMLA laserwriter (Heidelberg Instruments). Similar to Ref.^22^, the design consists of a 5 µm-tall and 440 µm-wide constriction area aligned with an adjacent 50 µm-tall perfusion channel for cell loading, chambers that end at the constriction area, and a bypass channel to equilibrate the fluid levels between the reservoirs positioned at the outer sides of the chamber.

The first layer of the design consisted of 5 µm SU-8 2005 photoresist (Kayaku Advanced Materials) and was spun on a 4-inch silicon wafer that was soft baked and post-baked at 95 °C for 2 minutes. The second layer consisted of 45 µm SU-8 3050 (Kayaku Advanced Materials) and was soft baked at 95 °C for 15 minutes, post-baked for 1 minute at 65 °C and 5 minutes at 95 °C, and developed with a SU-8 developer (Sigma Aldrich). After developing, trichloro(1H,1H,2H,2H-perfluoroctyl)silane (Sigma Aldrich) was coated on the master mold. Microfluidic chips were made from PDMS (Sylgard 184, Dow Corning), prepared with a curing agent with a 10:1 (w/w) ratio. The PDMS was poured on the silicon mold, degassed and cured at 65 °C for 3 hours. Reservoirs and cell-loading ports were punched with a revolving punch plier (Knipex) and a 0.75 mm diameter punch (Rapid-core, Well-tech), respectively. PDMS chips and glass coverslips were plasma cleaned (Harrick Plasma) at 30 W for 150 seconds and bonded overnight at 65 °C.

### Microfluidic migration assays

For live-cell migration experiments, microfluidic migration chips were sterilized with 70% ethanol and subsequently washed three times with MQ and once with PBS. Chips were coated with collagen by adding 100 µL 100 µg/mL PureCol type 1 bovine collagen (Advanced Biomatrix) in PBS through the cell-loading ports and incubating for 2 hours at room temperature. Chips were washed thrice with PBS and once with cell culture medium. Next, 6 µL containing a total of 30,000 cells was added to each chip via the cell-loading ports. The reservoirs were filled with cell culture media and the chips were incubated overnight at 37°C and 5% CO_2_ in a 10 cm culture dish, together with a 15 mL falcon tube cap filled with MQ next to the device to prevent evaporation. One hour before imaging, cell culture medium was removed from the reservoirs and replaced with cell culture medium containing live-cell dyes: Hoechst 33342 (Thermo Fisher, 1:10,000 dilution) for staining cell nuclei and Cytotracker Orange (Thermo Fisher, 1:1000 dilution) for staining the cell cytoplasm. After one hour incubation at 37 °, the device was sealed with a glass coverslip.

Live-cell imaging of cell migration through the constrictions was imaged on a Stellaris 8 confocal microscope (Leica), equipped with a supercontinuum white light laser, 405 nm laser and three hybrid detectors. Imaging was performed with the 405 laser, a 488 nm laser line and a 20x/0.75 air objective. Time-lapse image series were acquires with time intervals of 15 minutes over a total period of 13-14 hours. Environmental control with an box incubator (Okolab, #158206046) regulated the temperature at a constant 37 °C and 5% CO_2_. To analyze migration from the time-lapse series, we drew regions of interest as rectangles in between the diameters of two adjacent pillars (circles). First, the number of cells within these regions was counted. We identified cells as ‘squeezing’ if the cell entered and left the constriction region within the observation time and squeezed through the constrictions. We identified cells as ‘stuck’ when cells spent 10 or more frames (> 135 minutes) in a constriction region without squeezing to the other side and did not squeeze through any constriction during the observation time. In addition, for squeezing cells we counted the number of frames the Cytotracker or Hoechst signal was within the constriction region, to determine the amount of time cells and nuclei spent within the constriction during squeezing. Because time-lapses were created with 15 minute intervals, we rounded to the nearest 15 minute sampling interval. To analyze the decision making of the cells migrating through the bifurcating constrictions, we manually counted which constriction the cells migrated in.

### Western Blot analysis of vimentin expression

MV3 cells and HT1080 cells were both seeded in 6-well plates (Thermo Fisher) with 300,000 cells/well and incubated overnight to ensure cell attachment. In the morning, cells were washed with PBS and lysed with 100 µL cold radioimmunoprecipitation buffer (RIPA, Thermo Fisher). Lysed samples were agitated at 4° C for 30 minutes and stored at -20 °C. Loading samples were made by adding Laemmli buffer (2x, Bio-rad) and 4% β-mercaptoethanol (Sigma Aldrich) to the lysed samples. Next, they were incubated at 95°C for 5 minutes. Sodium dodecyl sulfate-polyacrylamide gel electrophoresis (SDS-PAGE) was performed with Mini-PROTEAN TGX gels (Bio-rad) using 100 V for approximately 1.5 hours. Western Blotting was executed with a Trans-Blot Turbo Transfer System (Bio-rad) and Trans-Blot Turbo Mini 0.2 µm PVDF Transfer Packs (Bio-rad). The membranes were blocked in 5% Bovine Serum Albumin (BSA, Thermo Fisher) in phosphate buffered saline (PBS, company) overnight on a shaker at 4 °C. Membranes were stained with primary antibodies (mouse antivimentin (1:2000, #ab8978, Abcam) and rabbit anti-GAPDH (#CST2118S, Bioke)) in 5% BSA overnight on a shaker at 4°C. Membranes were washed thrice with 0.1% Tween (Sigma Aldrich) in PBS (PBS-T) on a shaker, and incubated for 3-5 hours with secondary antibodies: rabbit anti-mouse HRP (#ab97051, Abcam) and goat anti-rabbit HRP (#ab6728, Abcam), 1:5000 in PBS-T. Afterwards, membranes were washed thrice with PBS-T and imaged with an enhanced luminol-based chemiluminescent substrate kit (Thermo Fisher) on a gel imager (Bio-rad).

### Statistical Analysis

Statistical analysis was performed using Microsoft Excel. Two-tailed Student’s t-tests were performed using the TTEST function. Statistical details of experiments are found in the figure legends and method details. P-value results from t-tests are indicated by: (ns) = p ≥0.05, (*) = p<0.05, (**) = p<0.01, (***) = p<0.001. Error bars represent the standard error of the mean.

See the supplementary material for the geometrical assumptions used to calculate theoretical strain, simulation of pressure distribution across the microfluidic cell deformability device, cell migration videos, and supplemental result figures references in the result and discussion sections.

## ACKNOWLEDGEMENTS

A.V.D.N., J.P.C. and G.H.K. gratefully acknowledge funding from the OCENW.GROOT.20t9.O22 project *The Active Matter Physics of Collective Metastasis* and the VI.C.182.004 project *How cytoskeletal crosstalk makes cells strong* (NWO Talent Programme), both financed by the Dutch Research Council (NWO). R.C.B. and P.E.B. gratefully acknowledge funding from the European Research Council (ERC) under the European Union’s Horizon 2020 research and innovation program (Grant agreement no. 819424).

## Ethics approval statement

Ethics approval is not required.

## Author contributions

**Anouk van der Net**: Conceptualization, Data curation, Formal analysis, Investigation, Methodology, Visualization, Writing - original draft. **Ruben C. Boot**: Conceptualization, Data curation, Formal analysis, Investigation, Methodology, Visualization, Writing - original draft. **Imke van Dijk**: Data curation, Formal analysis, Investigation, Methodology, Visualization. **James P. Conboy**: Investigation, Methodology, Formal analysis. **Pouyan E. Boukany**: Conceptualization, Funding acquisition, Visualization, Writing - original draft. **Gijsje H. Koenderink**: Conceptualization, Funding acquisition, Visualization, Writing - original draft.

## DATA AVAILABILITY STATEMENT

The data that support the findings of this study are available from the corresponding authors upon reasonable request.

