## Supplemental file for "Cancer cell deformability impacts the rate of confined migration but not decision making"

### SUPPLEMENTARY INFORMATION

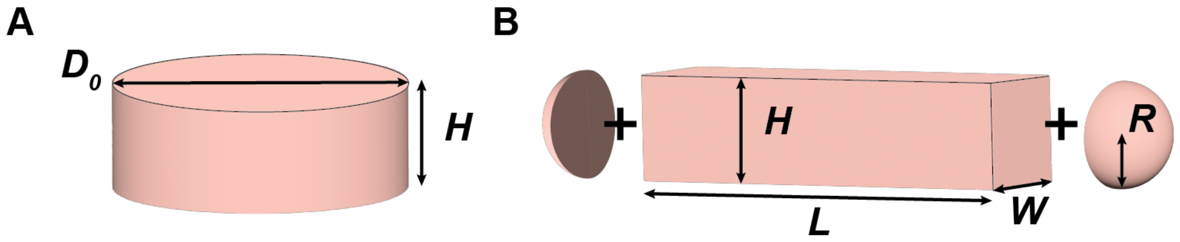


FIG. S1. Calculation of theoretical strain for cells forced through constrictions in the limit of volume conservation. Geometrical assumptions made to derive the theoretical strain $S_*$ in case of cell volume conservation when transiting through a narrow constriction with a square cross-section. (A) The cell being geometrically confined into a cylindrical shape at the inlet, with diameter $D_0$ and height $H$. (B) The cell being geometrically confined inside the constriction channel: the volume of the cell was determined as the sum of a beam of length $L$ with width $W$ and height $H$ and two hemispheres with a radius $R$ (= $\frac{1}{2}W = \frac{1}{2}H$).

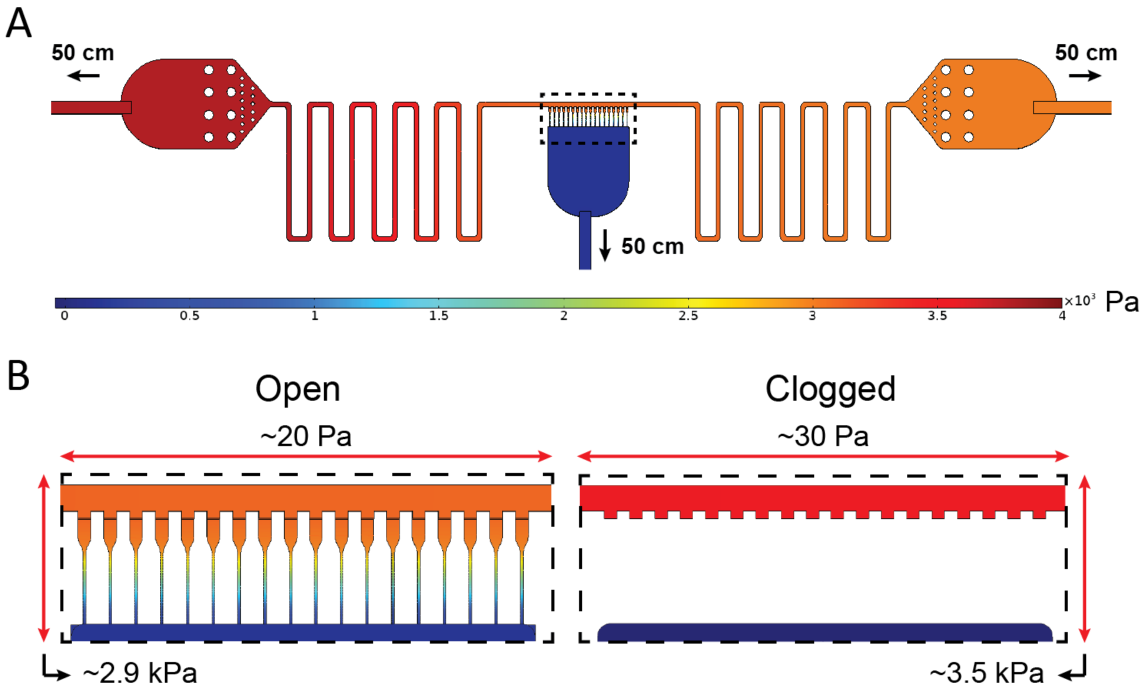


FIG. S2. Simulation of pressure distribution across the microfluidic cell deformability device. (A) 3D numerical simulation of the pressure distribution assuming the three ports are connected to 50 cm long rectangular channels mimicking the experimental tubing. A pressure gradient of 1 kPa is simulated between the far end of the tube connected to the cell entry port $P_1$ (4 kPa) and the tube connected to the perfusion port $P_2$ (3 kPa), while the far end of the tube connected to the outlet port $P_3$ is kept at atmospheric pressure (0 kPa). (B) Zoomed-in image showing a simulation of the horizontal and vertical pressure gradients across the microchannel array, demonstrated by the dashed rectangle in (A), for two different configurations: all microchannels are open (left) and all microchannels are clogged (right).

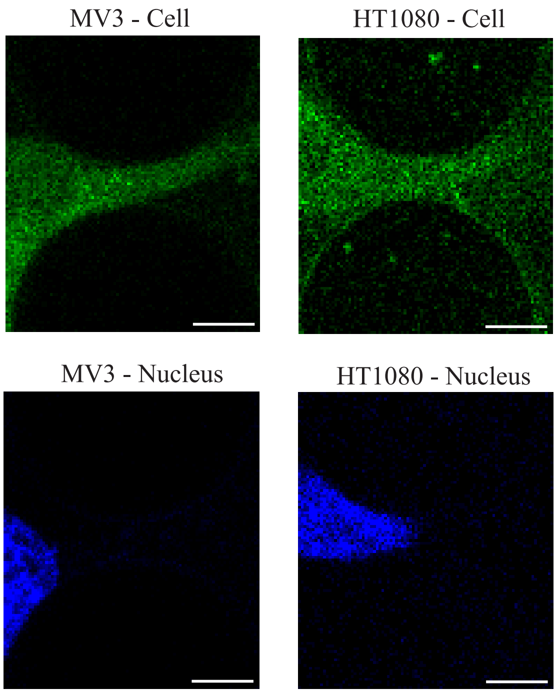


FIG. S3. Video stills for cell squeezing videos of HT1080 and MV3 cells. Scale bars are 15 μm. Multimedia available online.

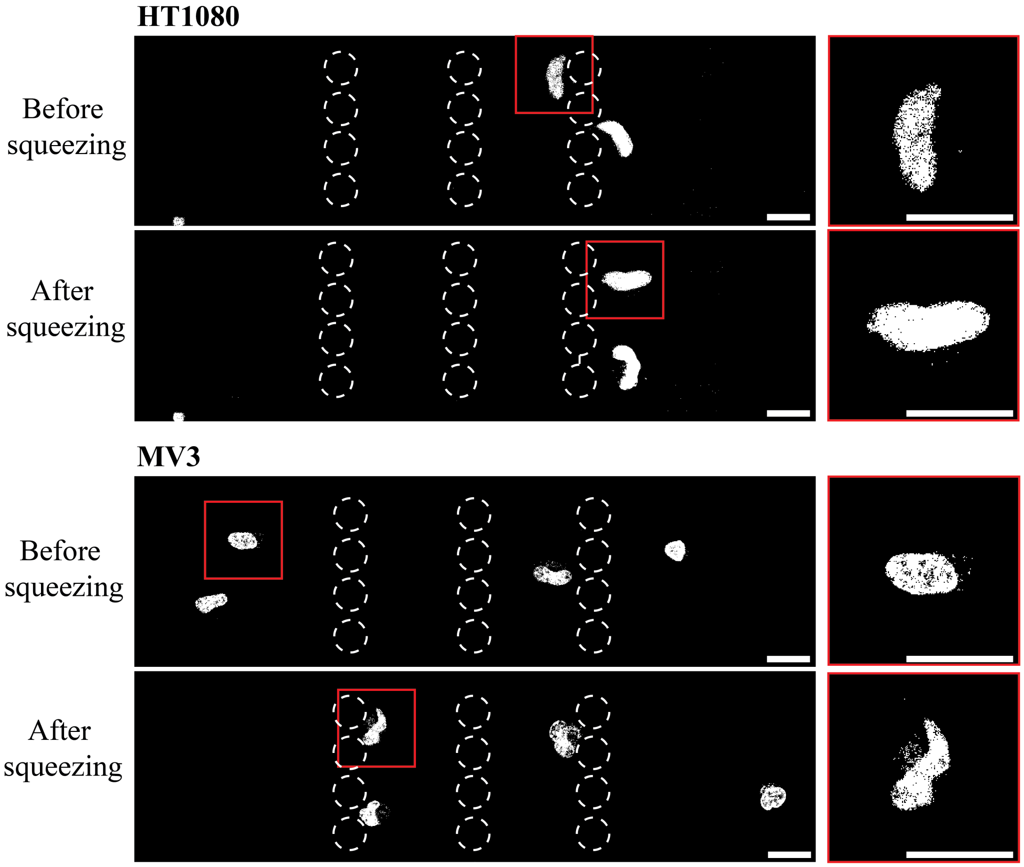


FIG. S4. Nuclear deformations after cell squeezing through microfluidic constrictions. Fluorescence images showing nuclear shapes of HT1080 and MV3 cells stained with Hoechst, before and after migrating through a constriction region. Pillars with constriction regions are indicated with white dashed circles. Zoomed-in views of nuclei indicated with red rectangles on the left are shown on the right. Scale bars are 25 µm.

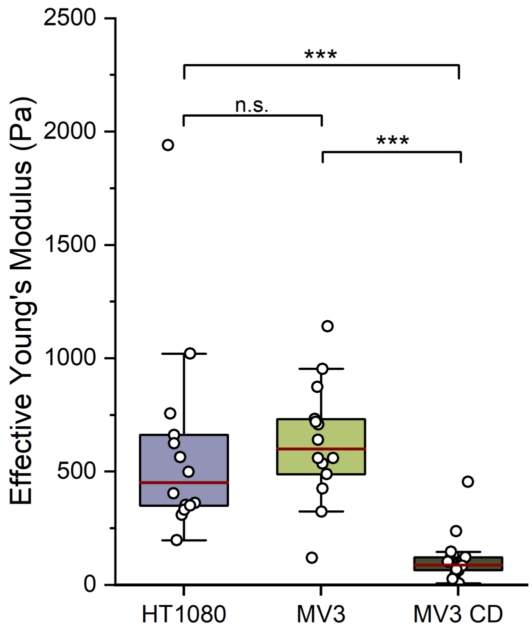


FIG. S5. Distribution of Young's moduli of adherent HT1080 cells, MV3 cells, and MV3 cells treated with cytochalasin D on glass, measured by microindentation using a probe with a spherical tip of radius 3 µm. Data are represented as box plots. The red lines represent the median values. (***) = p < 0.001 and (n.s.) = nonsignificant. n=14 for all conditions.

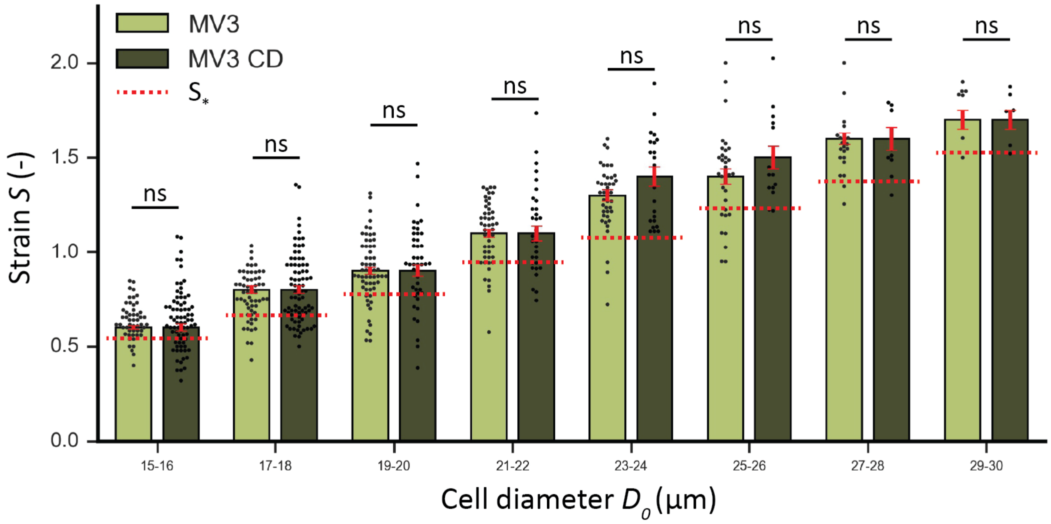


FIG. S6. Strain measurements for MV3 cells in response to Cytochalasin D treatment using the microfluidic cell deformability device. Histograms comparing the strain $S$ for control MV3 cells (light green) and MV3 cells treated with cytochalasin D (dark green), binned for cell diameter $D_0$. Red dotted lines depict the theoretical strain $S_*$ in the limit of volume conservation, calculated using Eq. (5) and inserting the largest $D_0$ for each bin (eg. $D_0$ = 16 µm for 15-16 bin. (ns) = nonsignificant. Error bars are SEM.

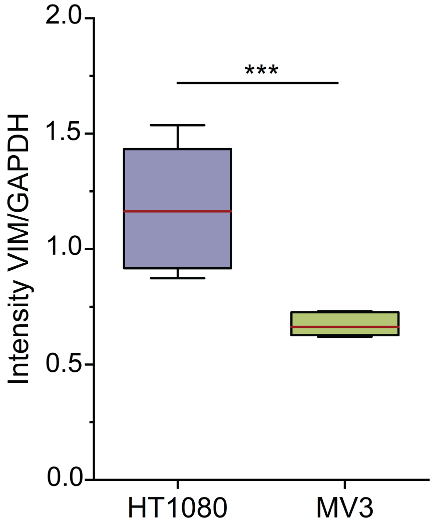


FIG. S7. Vimentin expression levels in HT1080 and MV3 cells. Boxplot showing intensities of the vimentin bands in a Western Blot for HT1080 (purple) and MV3 (green) cells, relative to GAPDH. The red lines indicate the median. (***) = $p < 0.001$. For each condition, n = two biological replicates, each subtracted from three different background spots, resulting in n = 6. Full blots are provided in Figure S9.

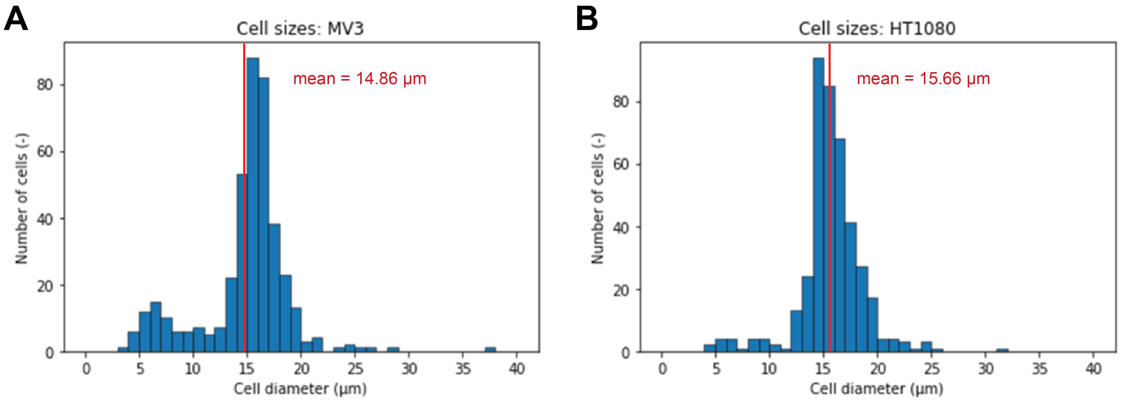


FIG. S8. Cell size distributions measured by an automated cell counter. Binned cell sizes for MV3 cells (A) and HT1080 cells (B). Bin size = 1 µm, n = 408 cells per cell line.

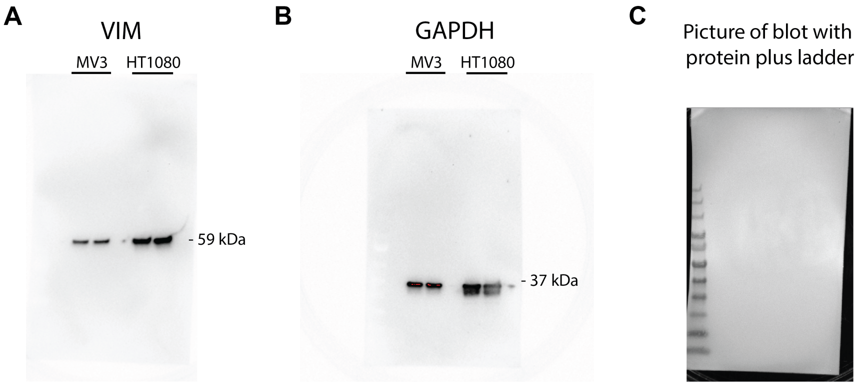


FIG. S9. Full size Western Blots for probing vimentin expression levels in HT1080 and MV3 cells. Western Blots for MV3 and HT1080 showing immunostained bands for (A) vimentin and (B) GAPDH, (C) and a picture of the blot showing the protein plus ladder.
